# Mechanical activation drives tenogenic differentiation of human mesenchymal stem cells in aligned dense collagen hydrogels

**DOI:** 10.1101/2021.11.25.470026

**Authors:** Hyeree Park, Showan N. Nazhat, Derek H. Rosenzweig

**Affiliations:** Department of Mining and Materials Engineering, McGill University, Canada; Division of Orthopedic Surgery, McGill University, Canada; Injury, Repair and Recovery Program, Research Institute of McGill University Health Centre, Canada

**Keywords:** gel aspiration-ejection, dense collagen, hydrogel, tendon, ligament, alignment, anisotropy, mechanobiology, mesenchymal stem cells

## Abstract

Tendons are force transmitting mechanosensitive tissues predominantly comprised of highly aligned collagen type I fibres. In this study, the recently introduced gel aspiration-ejection method was used to rapidly fabricate aligned dense collagen (ADC) hydrogel scaffolds. ADCs provide a biomimetic environment compared to traditional collagen hydrogels that are mechanically unstable and comprised of randomly oriented fibrils. The ADC scaffolds were shown to be anisotropic with comparable stiffness to immature tendons. Furthermore, the application of static and cyclic uniaxial loading, short-term (48 h) and high-strain (20%), resulted in a 3-fold increase in both the ultimate tensile strength and modulus of ADCs. Similar mechanical activation of human mesenchymal stem cell (MSC) seeded ADCs in serum- and growth factor-free medium induced their tenogenic differentiation. Both static and cyclic loading profiles resulted in a greater than 12-fold increase in scleraxis gene expression and either suppressed or maintained osteogenic and chondrogenic expressions. Following the 48 h mechanoactivation period, the MSC-seeded scaffolds were matured by tethering in basal medium without further external mechanical stimulation for 19 days, altogether making up 21 days of culture. Extensive cell-induced matrix remodeling and deposition of collagens type I and III, tenascin-C and tenomodulin were observed, where initial cyclic loading induced significantly higher tenomodulin protein content. Moreover, the initial short-term mechanical stimulation elongated and polarized seeded MSCs and overall cell alignment was significantly increased in those under static loading. These findings indicate the regenerative potential of the ADC scaffolds for short-term mechanoactivated tenogenic differentiation, which were achieved even in the absence of serum and growth factors that may potentially increase clinical translatability.

**Graphical abstract:** 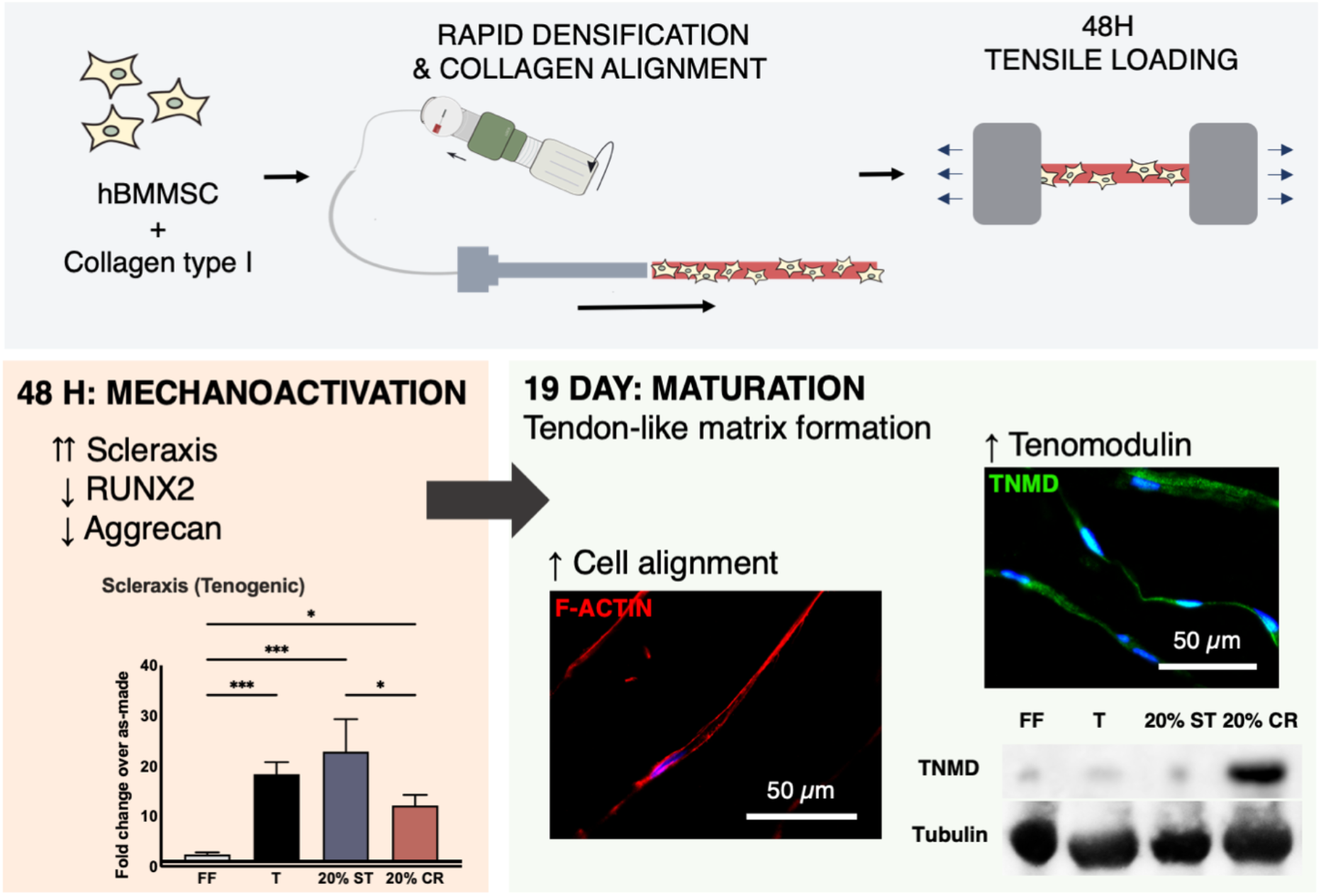

## 1. Introduction

Tendons and ligaments are mechanosensitive tissues that enable the load-bearing and force-transmitting functions of the body. Their injuries account for approximately 50% of the 33 million musculoskeletal injuries reported in the United States alone [1], costing approximately $30 billion per year [2]. These tissues have limited self-healing capacities due to lack of vasculature and reduced mechanical stimuli necessary for full regeneration. Autografts are the current gold standard, but re-rupture rates after reconstruction surgeries can be as high as 11.9% and 29% in anterior cruciate ligament and rotator cuff tendon, respectively [3,4], and often requiring further surgeries. Tendon tissue engineering aims to provide alternatives to autografts by providing a biocompatible material that acts as a scaffold for cell reprogramming and tissue remodeling *in vitro* or *in vivo* to fabricate a tendon-like tissue. Mesenchymal stem cells (MSCs) are multipotent and self-renewing cells that can be isolated from various tissues including bone marrow, adipose and tendons. These cells have shown great potential within regenerative medicine, where MSC differentiation has predominantly been achieved by exposure to growth factors. However, although tenogenic differentiation of MSCs *in vitro* has been achieved through supplementing growth factors in culture medium, such as transforming growth factor-β and fibroblast growth factor-2, a consensus on optimal dosages is yet to be determined [5–9]. Moreover, the exact pathways that are activated for tenogenesis remains unclear.

In lieu of growth factors, the mechanical microenvironment provided by the scaffold matrix has been shown to influence MSC differentiation into various lineages including tenogenic, osteogenic and chondrogenic lineages, indicating their mechanoresponsive malleability [10–14]. Even without external mechanical stimuli, the matrix stiffness can influence MSC differentiation [15–19]. For example, it has been shown that MSCs undergo either neurogenic, myogenic, or osteogenic differentiation when seeded in 2D on polyacrylamide gels of increasing stiffness [15]. Scaffold stiffness similar to embryonic tendon has also been observed to promote the tenogenic differentiation of seeded MSCs [7,20,21]. Since tendons are composed of 60 – 85% (dry weight) collagens, of which 95% is type I [22,23], *in vitro* reconstituted collagen type I hydrogels are commonly investigated as a 3D tendon scaffold. However, extracted self-assembled collagen hydrogels, exhibit poor mechanical properties and random fibrillar orientations [24,25]. These properties are not ideal as tendon scaffolds as they do not reflect the structure and anisotropy of the load-bearing aligned tissue and are mechanically weak.

Recently, the gel aspiration-ejection (GAE) method has been introduced, which rapidly densifies and remodels collagen hydrogels into aligned dense collagen (ADC) systems (Figure 1A) [26–28]. This process increases the collagen fibrillar density (CFD, or collagen content) through the removal of the casting fluid, and the applied pressure differential increases the alignment of the resulting hydrogel [26,27]. It has been shown that the modulus of ADCs can reach 3 MPa [26], thereby approaching those of immature bovine cruciate ligament and neonatal chick tendons [29,30]. Moreover, the GAE method also imposes the reorientation of 3D seeded cells in tandem with the directionality of the compacted collagen fibrils [26–28]. This indicates that ADCs provide a microenvironment resembling the composition and structure of tissue with highly aligned collagens. Due to the composition and anisotropy exhibited, we hypothesized that ADC hydrogels may be an appropriate scaffold for tendon tissue regeneration.

**Figure 1.**
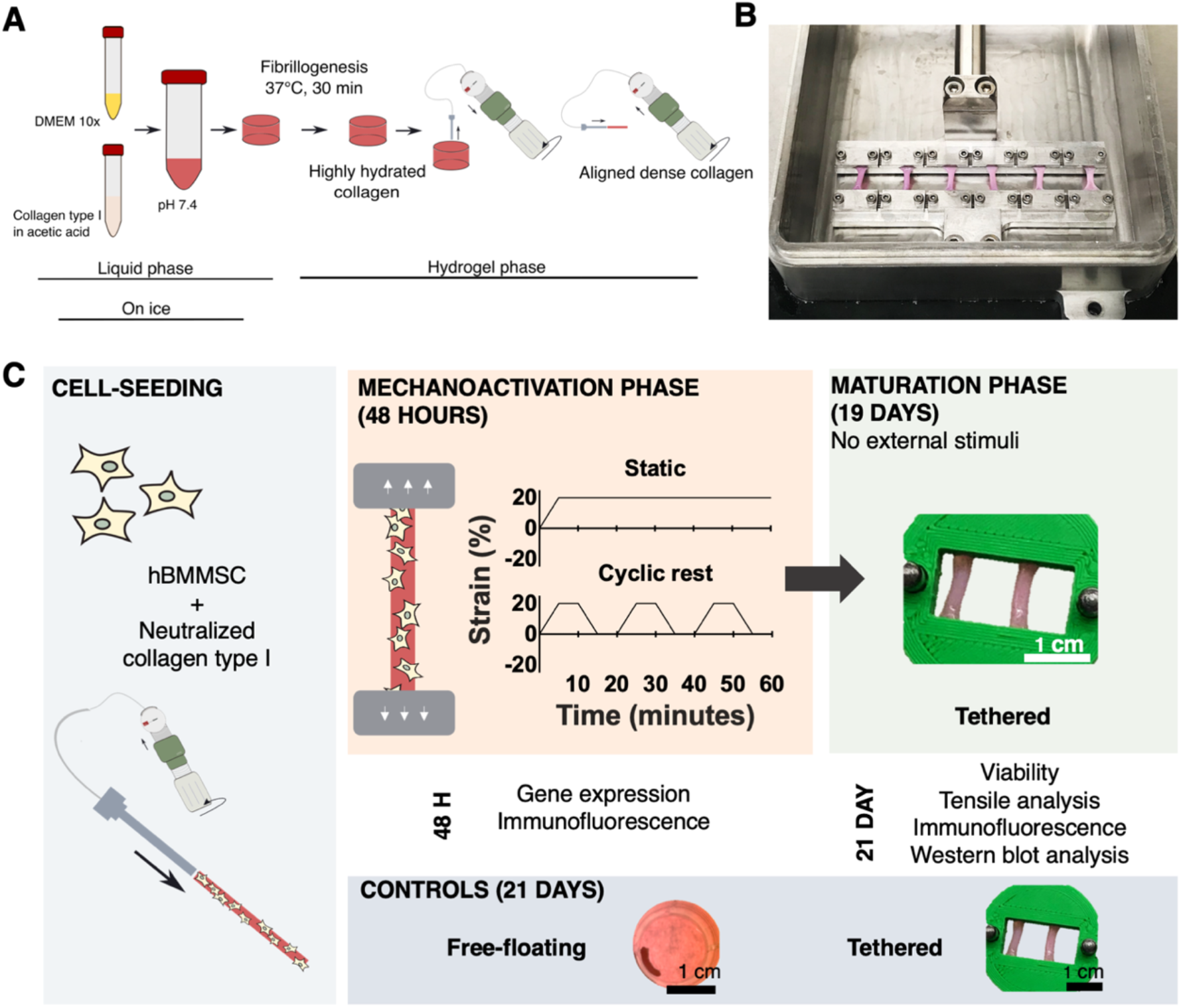
(A) GAE method and fabrication of ADC gels. (B) Loaded bioreactor for uniaxial stimulation. (C) Outline of hBMMSC-seeded culture and analyses. ADCs are seeded with hBMMSCs in its liquid phase, resulting in 3D embedded cells within the scaffold. A 48 h mechanoactivation phase was performed, using either 20% static strain or 20% cyclic rest profiles. These scaffolds subsequently underwent a maturation phase for 19 days by tethering in a mould without external mechanical stimulation. Free-floating and tethered ADCs were cultured for 48 h and 21 days to act as controls.

Although tenogenic differentiation of MSCs in both 2D and 3D culture *in vitro* has been achieved predominantly through growth factors [31–34], it can also be driven by external mechanical stimulation [10–14]. When mechanical stimuli have been applied for tenogenic differentiation, uniaxial cyclic loading between 3 – 10% strain has been used to mimic physiological strains and minimize overloading (>9% strain), which can cause apoptosis in *ex vivo* tendons and tendon progenitor cells in 2D [35]. However, MSCs may be more resilient as they have been shown to be viable following up to 20% cyclic loading [36] and may respond differently to mechanical stimulation than tendon derived cells. For instance, 3 – 10% cyclic straining has been shown to induce osteogenesis in MSCs [37–39], suggesting the need for higher strain levels to induce tenogenesis. In addition, the 3D seeding of the cells may also provide resistance to higher strain levels.

Compliance of 3D scaffolds with high strains may be limited depending on scaffold mechanical properties, especially when matched to physiological stiffness and ultimate strains. Previous investigation of randomly oriented dense collagen hydrogels of comparable CFD to ADCs have shown increased mechanical properties when stimulated acellularly at 20% strain [40]. Additionally, MSCs were observed to maintain their tenogenic gene profiles after 48 h of cyclic loading and maturation of a further 48 h in the absence of further external mechanical stimuli [10]. Therefore, it was hypothesized that ADCs may be compliant with 20% uniaxial strain in this time frame, where seeded MSCs would undergo mechanoactivation followed by tenogenic differentiation through a maturation phase. We aimed to observe the tenogenic differentiation potential of mechanical stimulation on MSCs seeded in ADC hydrogel scaffolds, and therefore growth factors were omitted throughout both stimulatory and maturation stages and serum was omitted during the stimulatory phase. Short-term (48 h) and high-strain (20%) static and cyclic tensile strains were applied, and the ADC hydrogel and seeded-MSC response was investigated. Further, the MSC-seeded ADCs were tethered in a mould and cultured to evaluate the effect of longer-term maturation within this mechanoactivated hydrogel environment.

## 2. Experimental methods

### 2.1. Cell culture

Human bone marrow mesenchymal stem cells (hBMMSCs; RoosterBio, USA) were purchased and cultured in RoosterNourish™-MSC media kit (RoosterBio, USA) to proliferate up to passage 1. The cells were further cultured in basal media, high-glucose Dulbecco’s modified eagle medium (DMEM; Sigma-Aldrich, Canada) supplemented with 10% fetal bovine serum (FBS; Gibco, Canada) and 1% penicillin/streptomycin (P/S, Gibco, Canada). Cells were passaged at 80% confluency and were used up to passage 4.

### 2.2. Production of acellular and hBMMSC-seeded aligned dense collagen scaffolds

ADC gels were fabricated using the GAE method (Figure 1A) [26]. For acellular ADCs, a 4:1 volume ratio of 2.05 mg/mL rat-tail derived collagen type I (First Link, Ltd., UK) and 10x DMEM (First Link Ltd., UK), respectively, was neutralized with 5M NaOH (Fischer Scientific, Canada). Aliquots of 1 mL of the neutralized solution were cast in 48-well plates (Sarstedt, Germany) and kept at 37 °C for 30 minutes to undergo fibrillogenesis to form collagen hydrogels. An angioplasty inflation device (B. Braun, Germany) attached to needles of 8 gauge (G), 10G, 12G and 14G was used to aspirate and eject the gels, fabricating a meso-scale anisotropic dense collagen gel. Fabrication of hBMMSC-seeded ADC gels used 8:1:1 ratio of collagen type I, 10x DMEM and cell laden basal DMEM, where the first two components were initially mixed and neutralized, and hBMMMSCs in basal DMEM were subsequently seeded at 200,000 cells/mL pre-densification.

### 2.3. Mechanoactivation and maturation of aligned dense collagen scaffolds

ADC gels fabricated by 12G needles were subject to a short-term (48 h) mechanoactivation phase. Both acellular and hBMMSC-seeded ADCs were mechanically stimulated using a uniaxially stimulation bioreactor (MechanoCulture T6, CellScale Biomaterials, Canada; Figure 1B). The protocols were chosen based on mechanical loading cycles applied to randomly oriented dense collagen in literature [40]. ADCs were subject to 48 hours of 20% static strain (20% SS), 5 minutes strain and hold; and 20% cyclic rest (20% CR), 5 minutes strain, 5 minutes hold, 5 minutes release to baseline, and 5 minutes hold (Figure 1C). Acellular ADCs were submerged in 1x Dulbecco’s phosphate buffer saline (PBS) solution (Gibco, Canada) and cellular scaffolds, in high-glucose DMEM and 1% P/S without serum in an incubator at 37 °C. Post mechanoactivation, hBMMSC-seeded ADCs were subject to a maturation phase (19 days), where the scaffolds were tethered in custom 3D-printed polylactic acid moulds to account for cellular contraction (stl files attached as supplementary information). The maturation phase was carried out in basal media supplemented with 50 μg/ml ascorbic acid (Sigma, Canada) to promote extracellular matrix formation whilst maintaining the differentiation potential of hBMMSCs [41]. As controls of cellular investigations throughout both mechanoactivation and maturation phases, free-floating and tethered ADCs were cultured for 48 h or 21 days.

### 2.4. Aligned dense collagen hydrogel characterization

Collagen content within the ADCs was measured as previously described [27,42]. Acellular ADC gels (n = 5) were weighed before and after freeze-drying (BenchTop K VirTis, USA) for over 10 hours. The percentage gel collagen content (measured through the collagen fibrillar density parameter, or CFD) was calculated by their dry-to-wet weight ratio. Tensile testing was carried out on ADCs (n = 5) using a Univert mechanical tester (CellScale Biomaterials, Canada) fitted with a 10 N load cell, at a displacement rate of 0.1 mm/s. Cross-sectional areas were assumed from the internal diameters of the needles, *i.e.* 3.43, 2.69, 2.16 and 1.60 mm for 8G, 10G, 12G and 14G, respectively. ADC gels were morphologically characterized by SEM. Samples were dehydrated by gradient ethanol baths followed by a 24-hour chemical dehydration in hexamethyldisilane (Sigma, Canada). The dehydrated samples were coated with 4 nm platinum using Leica Microsystems EM ACE600 (Leica, Austria). SEM images were obtained using FEI Inspect F-50 FE-SEM (FEI Company, USA) at 5kV and analyzed using Fiji (NIH, USA) to assess fibril thickness and directionality. Fibril thickness was manually measured over 50 measurements per image for three images captured per sample. Fibril orientation analysis was carried out using the directionality plug-in, where the ‘local gradient orientation’ algorithm was chosen [26]. The plug-in generated a histogram of fiber orientation fitted with a dominant Gaussian distribution [43].

### 2.5. Cell viability and number

LIVE/DEAD™ Viability/Cytotoxicity Kit (ThermoFisher, Canada) was used according to manufacturer’s instructions. Three images of three independent experiments (n = 3) of as-made scaffolds as well as those cultured to day 21 were captured using EVOS M5000 (ThermoFisher, Canada), and were quantified using the cell counter plug-in on ImageJ (NIH, USA). Cell number was quantified using the Hoechst 33258 DNA assay as previously described [44–46]. After 21 days of culture, scaffolds (n = 3) were submerged into 500 μL of 4 M guanidine hydrochloride (GuHCl, Sigma-Aldrich, Canada) buffer supplemented with a protease inhibitor (Roche Applied Science, USA) and gently agitated for 48 h at 4 °C. 1 μg/mL Hoechst 233258 (Invitrogen, USA) and serial dilutions of calf-thymus DNA (Invitrogen, USA) supplemented with 4M GuHCl were prepared. Supernatants and diluted calf-thymus DNA and were placed into 96-well plates and assessed using a Tecan M200, at 360 nm excitation and 460 nm emission. Standard curves were inferred from readings of the calf-thymus DNA, and the weight of DNA from a single cell was estimated to be 7 picograms [45].

### 2.6. Gene expression

500 μg of RNA was extracted from hBMMSC-seeded ADC gels (n = 3) using TRIzol reagent (Invitrogen, USA) as per manufacturer’s instructions following the initial 48 h of mechanical stimulation. Extracted RNA was reverse-transcribed using qScript cDNA synthesis kit (Quanta Biosciences, USA) and quantitative real time polymerase chain reaction (qRT-PCR) was performed using PowerUp™ SYBR™ Green Master Mix (Applied Biosystems, USA) with a StepOnePlus Real Time PCR System (Applied Biosystems, USA). Scleraxis, runt-related transcription factors (RUNX2) and aggrecan were quantified and normalized to the housekeeping gene glyceraldehyde 3-phosphate dehydrogenase (GAPDH). Primer sequences can be found in Table 1. The average fold change in gene expression of experimental samples compared with as-made ADC controls was calculated using the 2^-ΔΔCt^ method [47].

**Table 1.**
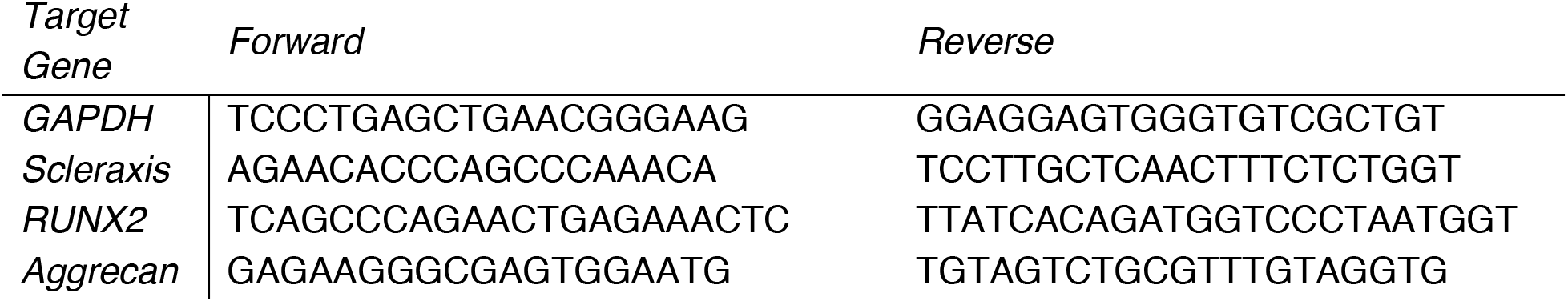
Primer sequences for qRT-PCR

### 2.7. Protein analysis

As made, 48 h and day 21 hBMMSC-seeded scaffolds (n=3) were fixed in 4% paraformaldehyde (Sigma, Canada) for 1 hour and submerged in gradient sucrose solutions from 10% to 30%. The fixed samples were embedded in O.C.T. compound (TissueTek, Canada), snap frozen at −80°C and cut to 7 *μ*m sections using CM1950 cryostat (Leica biosystems, Germany). Sections were permeabilized and blocked (5% BSA, 0.1% Triton-X 100 in PBS) for 45 minutes and incubated overnight at 4°C with one or two primary antibodies from different hosts. Primary antibodies used and their concentrations were as follows: alpha smooth muscle actin (ab5694 1:200, Abcam, Canada), collagen I (1:200, ab34710, Abcam, Canada), collagen III (1:200, ab7778, Abcam, Canada), collagen VI (1:10, 5C6, DSH, USA), tenascin-C (1:200, ab3970, Abcam, Canada), scleraxis (1:200, ab58655, Abcam, Canada) and tenomodulin (1:200, ab203676, Abcam, Canada). These were further incubated for 1.5 h in room temperature with AlexaFluor 488 donkey anti-rabbit IgG (1:250, A21206, Invitrogen, Canada) and/or AlexaFluor 555 Goat anti-Mouse IgG (1:250, A21422, Invitrogen, Canada). Where F-actin was co-stained, the secondary antibody was supplemented with Texas Red™-X Phalloidin (1:400, T7471, Invitrogen, Canada). Cellular alignment was determined by 10x F-actin staining (n = 3), where ImageJ (NIH, USA) and its directionality plug-in was used as described above with collagen fibrillar alignment.

Using the remaining supernatant of the GuHCl treated scaffolds, the supernatant was precipitated by anhydrous ethanol [45]. The protein concentration was measured using the BCA assay. 10 μg of extracted proteins (n = 3) were solubilized in 1 × NuPage® LDS Sample buffer (Invitrogen, USA) and was run in a 15 well 12% Tris-Gly precast gel (Invitrogen, USA) at 25 mA for 1 h and transferred onto an Amersham™ Protran® 0.2 μm nitrocellulose blotting membrane (GE Healthcare, Canada) at 300 mA for 1.5 h. The membrane was incubated with primary antibody overnight at 4°C with collagen I (1:1000 dilution, ab34710, Abcam, Canada), tenomodulin (1:1000, ab203676, Abcam, Canada) or tubulin (1:1000, ab7291, Abcam, Canada). The membranes were subsequently incubated with horseradish peroxidase (HRP) conjugated secondary antibodies for 1 h at room temperature, with goat anti-rabbit secondary antibody (1:2000, ab6721, Abcam, Canada) for tenomodulin and goat anti-mouse (1:5000, ab6789, Abcam, Canada) for tubulin. The immunoreactive bands were detected using Western Lightning™ Chemiluminescence Reagent Plus (Perkin Elmer, USA) and LAS 4000 Image Quant system (GE Healthcare, Canada). The sizes of the bands were analyzed using ImageJ software using the gel analysis plug-in.

### 2.8. Statistical analysis

Experimental data was analysed using Graphpad Prism™ (Graphpad Prism Software, USA.) Groups were analysed with an unpaired Student t-test, and where applicable, two-way ANOVA with multiple comparison (Turkey’s) was analyzed. Linear regression analysis was applied to correlate mechanical properties to the CFD.

## 3. Results

### 3.1.1. Characterization of acellular ADC hydrogels

ADC hydrogels were fabricated using the GAE method and needles between 8 and 14 G. The SEM micrographs of ADC surface morphology showed that the collagen fibrils were more aligned with a decrease in needle cross-sectional diameter (Figure 2A). The quantified alignment confirmed the relationship between the needle internal diameter and amount of alignment (Figure 2B-C), although the amount of alignment between 8 and 10G (p = 0.09) as well as 12 and 14G (p = 0.41) did not show significance. Another parameter that was correlated with the needle size was CFD (Figure 2D), *i.e.,* 5.4 ± 0.3 wt.% (8G), 5.8 ± 1.1 wt.% (10G), 7.9 ± 1.5 wt.% (12G) and 12.8 wt.% (14G). All ADCs were observed to increase in collagen content compared to the initial collagen concentration of 2.05 mg/mL, or approximately 0.2 wt.% CFD. As with the amount of alignment, the CFD of adjacent 8 and 10G hydrogels were not significant (p = 0.42).

**Figure 2.**
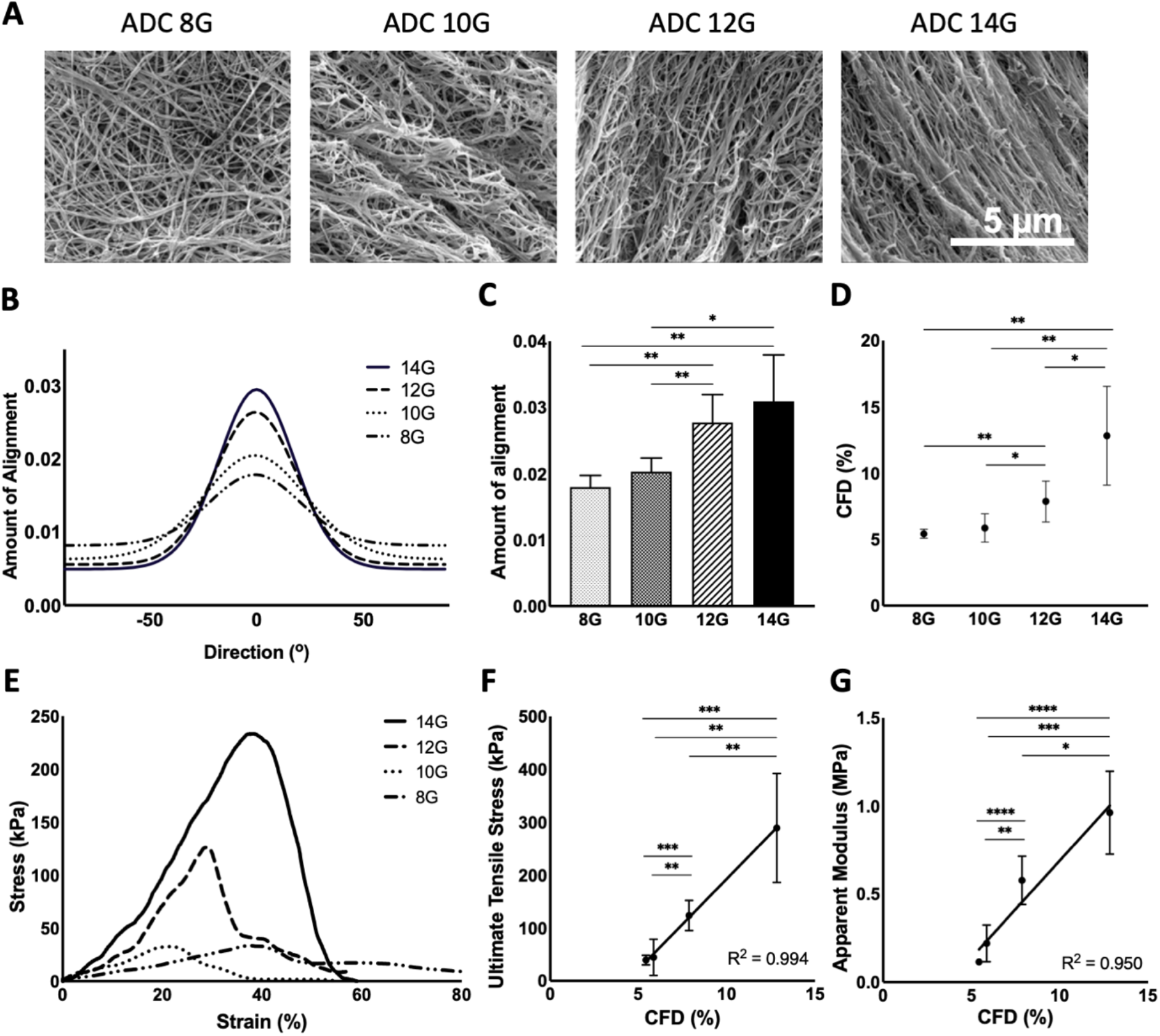
Characterization of as-made 8 – 14G acellular ADC gels. (A) SEM images of 8 – 14G ADC gels, scale bar = 5 *μ*m and (B) average directionality histogram and (C) amount of fibrillar alignment as depicted in SEM images. (D) Collagen fibrillar density (CFD) of the gels. (E) Representative tensile stress-strain curves for the 8 – 14G ADC gels and their (F) ultimate tensile stress and (G) apparent modulus, both fitted with a linear regression curve. Error bars ± SD and *p < 0.05, **p < 0.01, ***p < 0.001, and **** p < 0.0001 as per Student t-test.

The CFD had a direct impact on the tensile properties and positive correlation was seen where R^2^ values for ultimate tensile strength (UTS) and apparent modulus (AM) values were above 0.95 (Figure 2E-G). The UTS ranged between 40 ± 10 kPa (8G) and 290 ± 100 kPa (14G), while the AM between 0.11 ± 0.01 MPa (8G) and 0.96 ± 0.24 MPa (14G). Both UTS and AM values obtained were significantly different between needle gauges (p > 0.05), except for between 8G and 10G.

Acellular 12G ADC hydrogels were mechanically stimulated uniaxially for 48 h under static and cyclic tensile straining profiles, *i.e.,* 20% static strain and cyclic rest. The stimulated hydrogels showed no signs of material failure due to these strain levels. Post-straining, an increase in collagen fibrillar thickness was observed (p < 0.05 for both conditions against unstimulated control) (Figure 3A,D), but there was no significant difference (p > 0.05) of the collagen fibrillar alignment (Figure 3B-C). Tensile mechanical testing (Figure 3E-G) revealed that 20% static strain significantly increased UTS and AM, both by 2.7-fold (340 kPa, p = 0.005) and 3-fold (1.72 MPa, p = 0.001), respectively, whereas 20% cyclic rest significantly increased AM only by 2.6-fold (1.43 MPa, p = 0.043).

**Figure 3.**
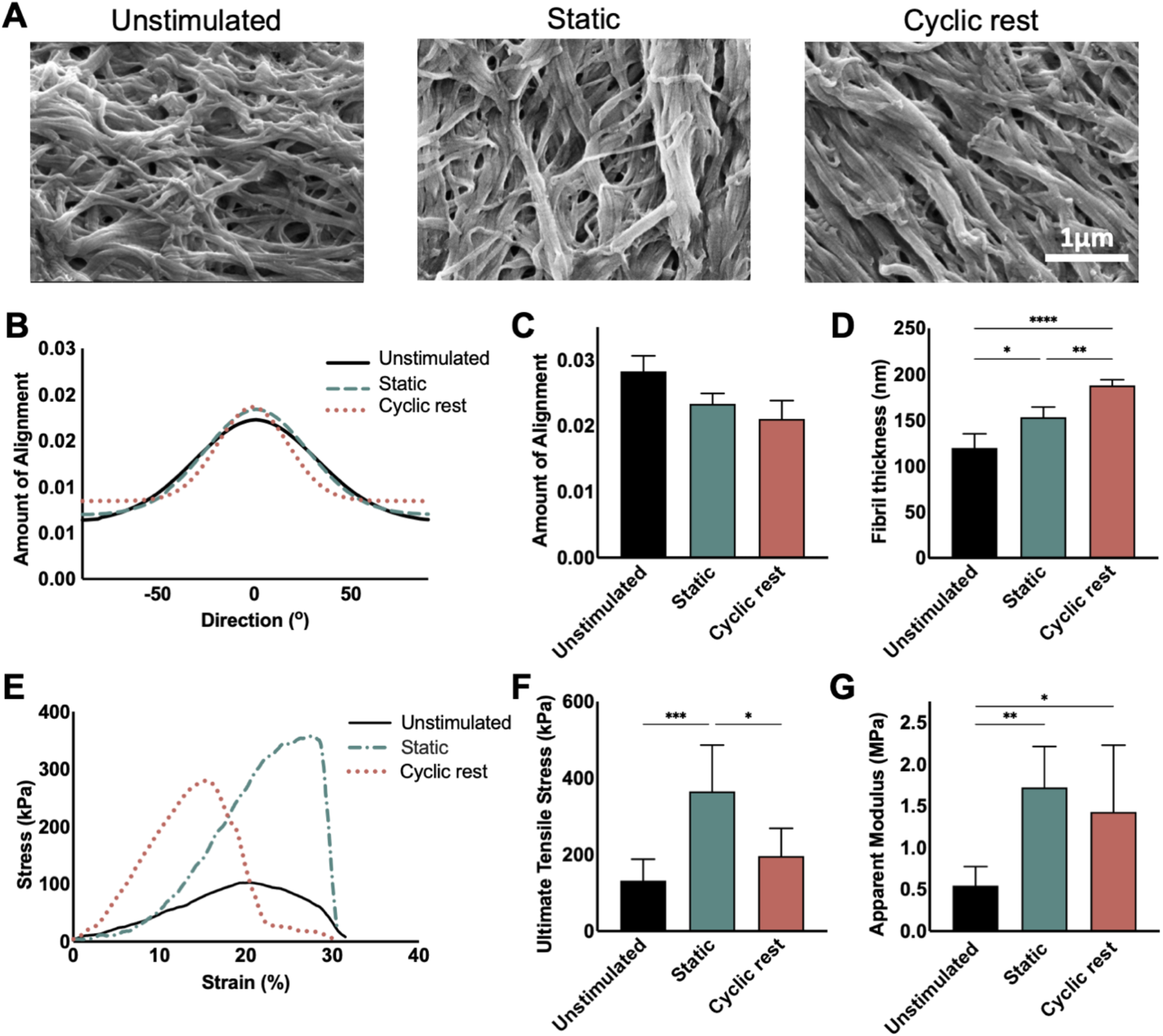
Characterization of acellular 12 G ADC gels after 48 h mechanical stimulation. (A) Representative SEM images of ADC gels post-stimulation, scale bar = 1 *μ*m. (B) Average directionality histogram, (C) amount of fiber alignment and (D) fibril thickness as depicted in SEM images. (E) Representative tensile stress-strain curves for the unstimulated and stimulated ADC gels and their (F) ultimate tensile stress and (G) apparent modulus. Error bars ± SD and *p < 0.05, **p < 0.01 and ***p < 0.001 as per student t-test.

### 3.1.2. Viability of cell-seeded ADCs

The 3D seeding of hBMMSCs in ADC gels was achieved pre-gelation and application of GAE, as previously reported [26–28,48]. The hBMMSC-seeded ADC gels were subjected to 20% static strain or 20% cyclic rest loading profiles for 48 h and kept tethered for subsequent 19 days, with controls that were free-floating and tethered for the 21-day culture (Figure 4). As made ADC gels showed 89 ± 2*%* viability and all cultured conditions showed between 70 to 83% viability at day 21 (Figure 4 A-C). Cellular proliferation throughout the 21-day culture was indicated by DNA quantification, where 2.5 and 3-fold increases, *i.e.,* between 500,000 and 600,000 cells, were observed from the initially seeded 200,000 cells per scaffold (Figure 4D). The ADCs provided a cytocompatible environment for the seeded hBMMSCs, where high viability post-fabrication and culture, as well as cell proliferation was observed.

**Figure 4.**
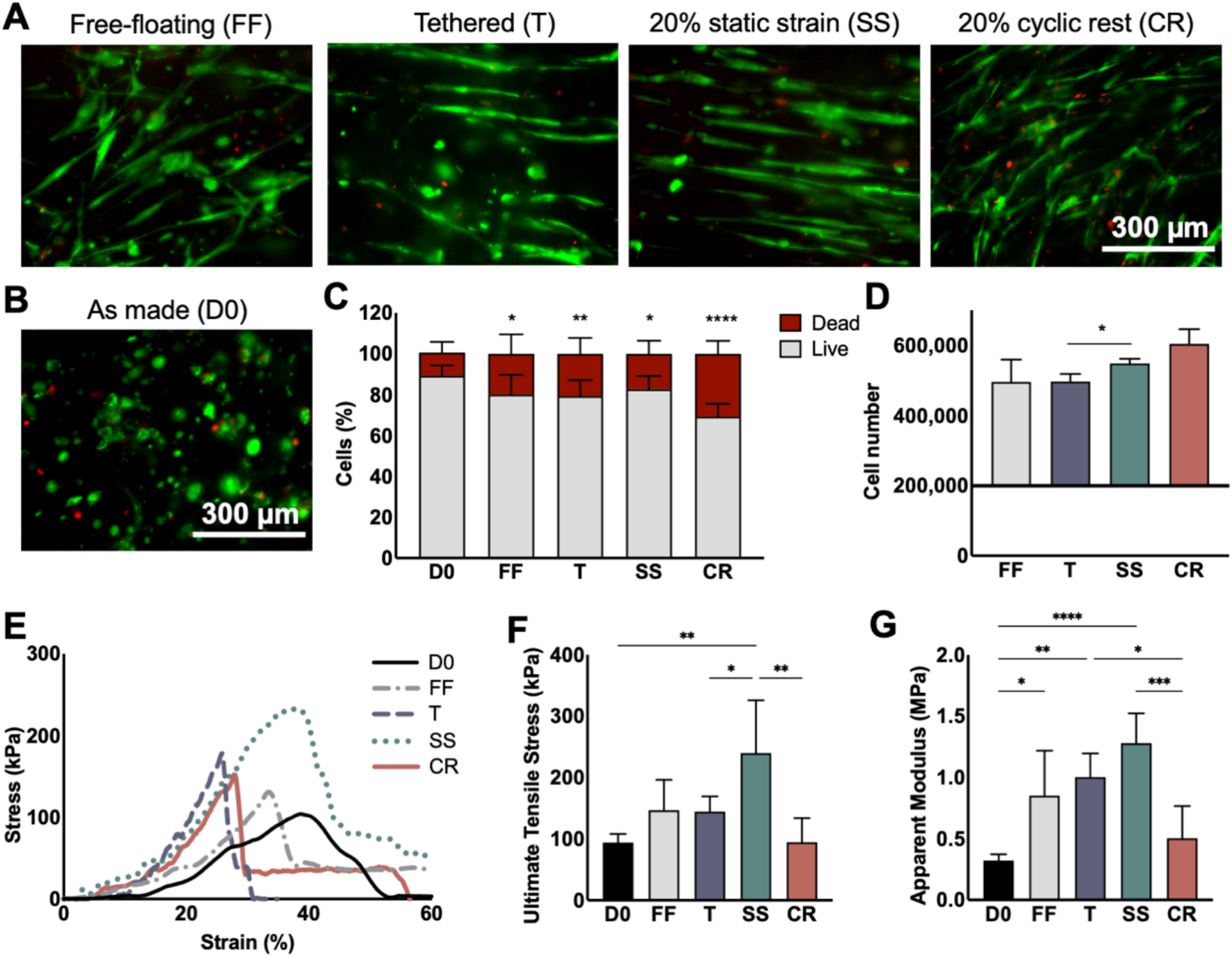
Viability and mechanical properties of hBMMSCs seeded in ADC gels cultured for 21 days (i.e., 19 days maturation post initial 48 h mechanostimulation). Representative Live/Dead images taken at 10x (A) at day 21 of free-floating (FF), tethered (T), 20% static strain (SS) and 20% cyclic rest (CR) and (B) as made at day 0 (D0). (C) Percentage live and dead cells from quantification of Live/Dead images. (D) The number of cells present at day 21 as quantified by the Hoechst 33258 DNA assay. (E) Representative tensile stress-stress curves at day 21 and their (F) ultimate tensile stress and (G) apparent modulus. Error bars ± SD and *p < 0.05, **p < 0.01, ***p < 0.001 and **** p<0.0001 as per Student t-test. Scale bars = 300*μ*m.

### 3.1.3. Tensile properties of cultured ADCs

The cellular as made ADC hydrogels did not show statistically significant mechanical properties compared to the acellular as made scaffolds (Figure 4E-G), suggesting that the 200,000 cells seeded per scaffold did not significantly affect the mechanical properties of the hydrogel (UTS p = 0.172 and AM p = 0.068). However, when cultured, the increase in mechanical properties indicated cell-induced remodeling (Figure 4E-G). Hydrogels exposed to 20% static strain showed significant increases from as made controls, reaching UTS of 240 ± 85 kPa (p = 0.006) and AM of 1.28 ± 0.24 MPa (p < 0.0001). These were also statistically significant with the UTS of tethered samples (p = 0.044) and cyclic rest (p = 0.009). In contrast, the cyclically loaded ADCs with the cyclic rest profile did not increase mechanical properties compared to the as made controls, with the UTS of 95 ± 39 kPa (p = 0.97) and AM of 0.502 ± 0.263 MPa (p = 0.17). Both cultured controls, free-floating and tethered hydrogels, significantly increased their mechanical properties from the cellular as made controls (p > 0.05) but were not statistically significant from each other.

### 3.1.4. Mechanoactivation drives tenogenic differentiation

The differentiation gene markers of the hBMMSCs seeded in the ADCs were investigated immediately after the 48 h mechanostimulatory period. The free-floating hydrogels trended inversely to the tethered and mechanically stimulated hydrogels. Scleraxis (Figure 5A.I), an early tenogenic marker [49–51], was upregulated for all loading profiles and controls. Tethered, static strain and cyclic rest ADCs increased scleraxis expression by over 12-fold, whereas free-floating hydrogels were upregulated by 2 ± 0.4-fold indicating a biophysical effect of the scaffold itself. Conversely, for the free-floating hydrogels, the osteogenic and chondrogenic markers, RUNX2 [52] and aggrecan [44] (Figure 5A.II, III), were upregulated by 11 ± 4 -fold and 9 ± 6 -fold, respectively, but were significantly suppressed or maintained at baseline for both loading conditions. RUNX2 expressions was downregulated by 5-fold (0.2 ± 0.07) for tethered ADCs and below detectable for static strain and cyclic rest groups. Aggrecan expressions were maintained for tethered and cyclic rest groups at 1.25-fold (0.8 ± 0.2) and 1.11-fold (0.9 ± 0.3), respectively, and suppressed below detectable for the static strain group.

**Figure 5.**
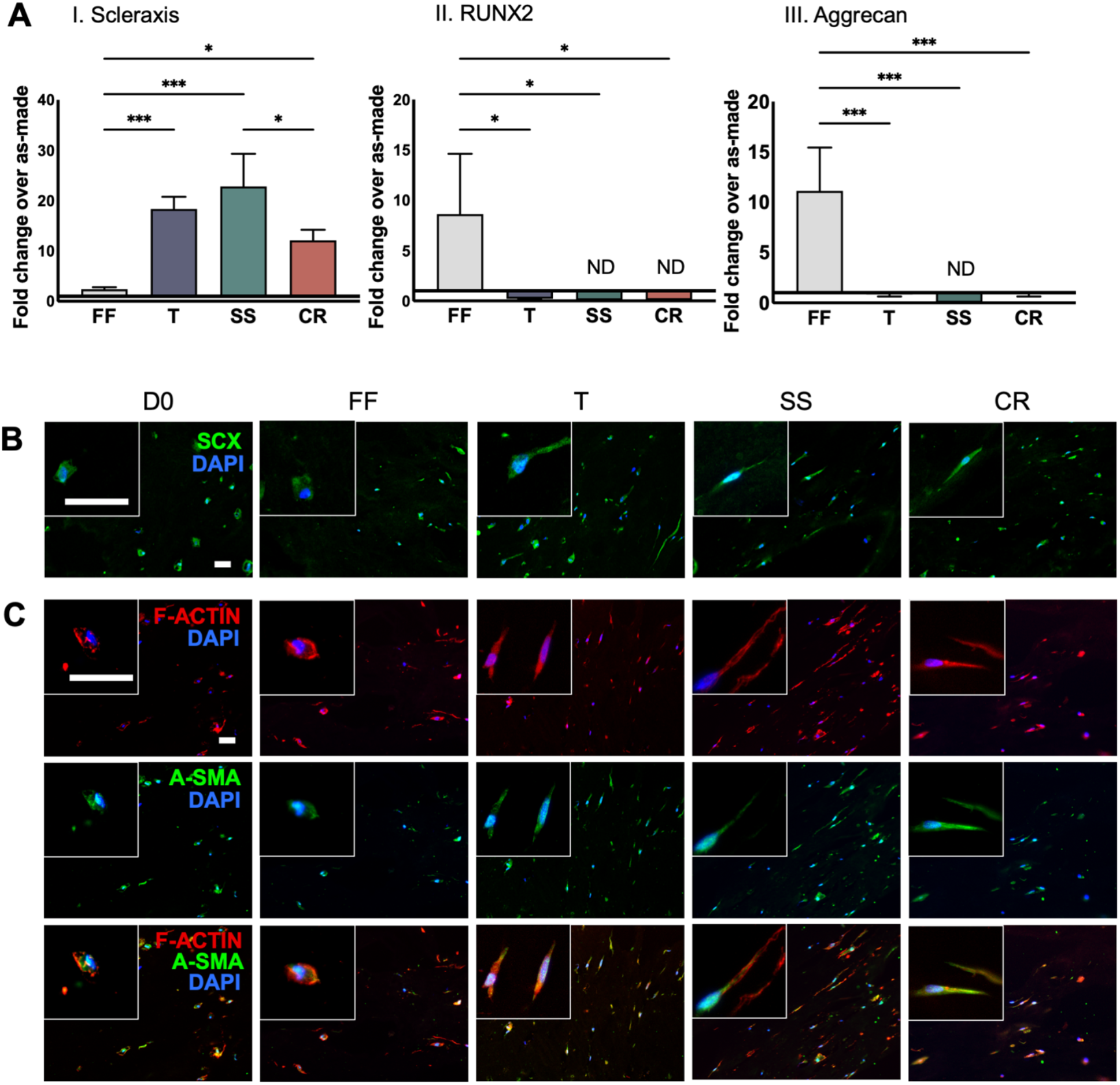
Characterization of hBMMSC-seeded ADCs after 48 hours of mechanoactivation through 20% static strain (SS) and 20% cyclic rest (CR) and their controls, free-floating (FF), tethered (T) and as made (D0). (A) Genetic markers of differentiation of uniaxially stimulated hydrogels, reported as fold change over as made (D0) controls: (I) Scleraxis (SCX), an early tenogenic transcription factor; (II) RUNX2, an early osteogenic marker; and (III) aggrecan (AGG) a chondrogenic marker. (B) Immunofluorescent staining of scleraxis (SCX). (C) Immunofluorescent staining of F-actin and alpha-smooth muscle actin (A-SMA) and their merged images. Error bars ± SD and *p < 0.05, **p < 0.01, ***p < 0.001 and ****p < 0.0001 as per two-way ANOVA post-hoc multiple comparison analysis. ND indicates not detected for fold change < 0.1. Scale bars = 50*μ*m.

The immunofluorescent staining of scleraxis at 48 h corroborated the scleraxis gene expressions, where it was detected in all conditions but more significantly in tethered and initially mechanically stimulated hydrogels (Figure 5B). The as made ADCs showed scleraxis stains, indicating that hBMMSCs inherently express scleraxis [53]. The morphology of the cells could be identified by the F-actin stains (Figure 5C). Immediately after seeding, the hBMMSCs were rounded, which was maintained by the cells in free-floating ADCs. In contrast, when tensile static or cyclic strains were applied, the cells became polarized and in the case of ADCs under static strain, the cells were also significantly elongated. All conditions showed perinuclear stains of alpha-smooth muscle actin (Figure 5C), an indicator for cellular contraction and matrix remodeling [54–56].

### 3.1.5. Tendon-like matrix formed during maturation

After the 48-h stimulatory period, the seeded ADC scaffolds were further cultured for 19 days, for a total of 21 days of differentiation and maturation. F-actin stain was used to both qualitatively observe cellular morphology and quantitatively investigate cellular alignment. High magnification images of F-actin (Figure 6A) indicated that the cells seeded in tethered, static strain and cyclic rest conditions were all polarized along the collagen fibrils showing elongation of the nucleus and cytoskeleton, thus exhibiting tendon-like morphology. The cellular elongation was dependent on the presence of initial tensile loading, as it was more histologically prominent for the static strain and cyclic rest hydrogels than those tethered. Conversely, cells in free-floating ADCs were more hypertrophic and the cytoskeleton filaments were not uniformly polarized as some filaments extended perpendicular to the collagen fibril orientation, which deviated from typical tenogenic cell morphology. Cellular alignment was quantified using low magnification (10x) F-actin images (Figure 6C-D), which revealed that holistically, the initially statically strained groups had the most aligned cells, which were significantly different from all other loading profiles (p < 0.05). All other groups were not significant in their amount of alignment. However, the morphological differences at the cellular level suggest that tensile straining is an important factor in the tenogenic differentiation of MSCs.

**Figure 6.**
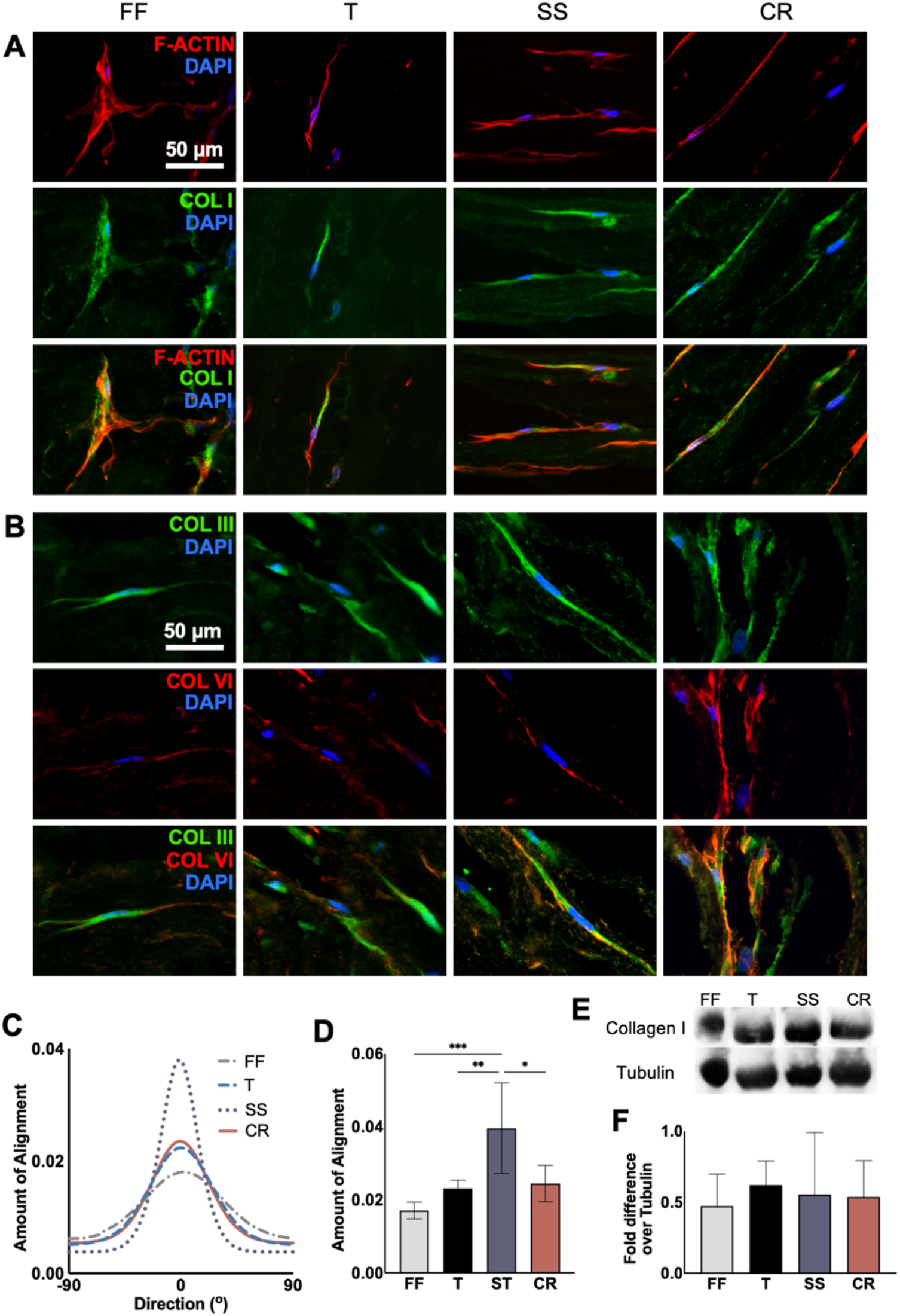
Cellular morphology and collagen formation of hBMMSC-seeded ADCs at day 21 after 48 h mechanoactivation and 19-day maturation of 20% static strain (SS) and 20% cyclic rest (CR) and their controls, free-floating (FF) and tethered (T). (A) F-Actin (red) and COL1 (green) with nucleus (blue) staining, (B) COL3 (green) and COL6 (red) with nucleus (blue) staining. (C) Average cellular alignment histograms quantified from 10x F-Actin staining and (D) the modes of the histograms. (E) Western blot protein analysis of COL1 in the ADC hydrogels and (F) densitometry analysis of COL1, where no treatment groups were statistically significant. Error bars ± SD and *p < 0.05, **p < 0.01 and ***p < 0.001 as per one-way ANOVA post-hoc multiple comparison analysis. Scale bars = 50 *μ*m.

Both fibril-forming collagen types I and III, and non-fibril-forming type VI were detected by immunofluorescent staining (Figure 6A-B). Collagen type I, the most prominent protein within the tendon extracellular matrix, was detected along with the F-actin filaments for all treatment groups. Western blot analysis identified non-crosslinked collagen type I (Figure 6E-F) where substantial levels of collagen type I formation was observed for all conditions. The densitometry was quantified and found to be non-significant between all treatment groups (p > 0.05). Collagen type III, the second most prominent protein within tendons, was also identified for all conditions. Where initial tensile loading was applied, it was expressed in a more elongated and interconnected manner. Collagen type VI is known to be present within various tissues including tendons, interacting with fibrillar collagens and proteoglycans, and is localized along the fibrillar matrix extending to the cell orientation [57]. The immunostaining showed the longitudinal extension of collagen type VI overlapping with the edge of the collagen type III stain, with higher levels of prominence within the static strain and cyclic rest profiles.

The tenogenic markers scleraxis, tenascin-C and tenomodulin were also investigated using immunofluorescent staining (Figure 7A-B), Scleraxis at day 21 was present for all conditions, but with prominent nucleic stains for those initially stimulated through static strain and cyclic rest. Tenascin-C is a glycoprotein that is found within tissues that undergo loading [58] and is commonly investigated as a tenogenic marker [49]. Where tensile loading was applied, by static strain and cyclic rest, Tenascin-C was identified in the nucleus as well as along the cytoskeleton and observed to be polarized and connected between cells. Tenomodulin, a mature tendon protein [59,60], was found to be most strongly positive in cyclic rest hydrogels, which was confirmed by the western blot analysis (Figure 7C-D), suggesting that cyclic loading is imperative to forming mature tendon-like matrix. The weak presence of Tenascin-C and tenomodulin in free-floating and tethered conditions indicate that the tensile loading may drive tenogenic differentiation of MSCs. The lack of tenomodulin detected in the western blot for static strain conditions support that cyclic tensile loading may be a key element in differentiating MSCs and stimulating their fabrication of a mature tendon-like matrix.

**Figure 7.**
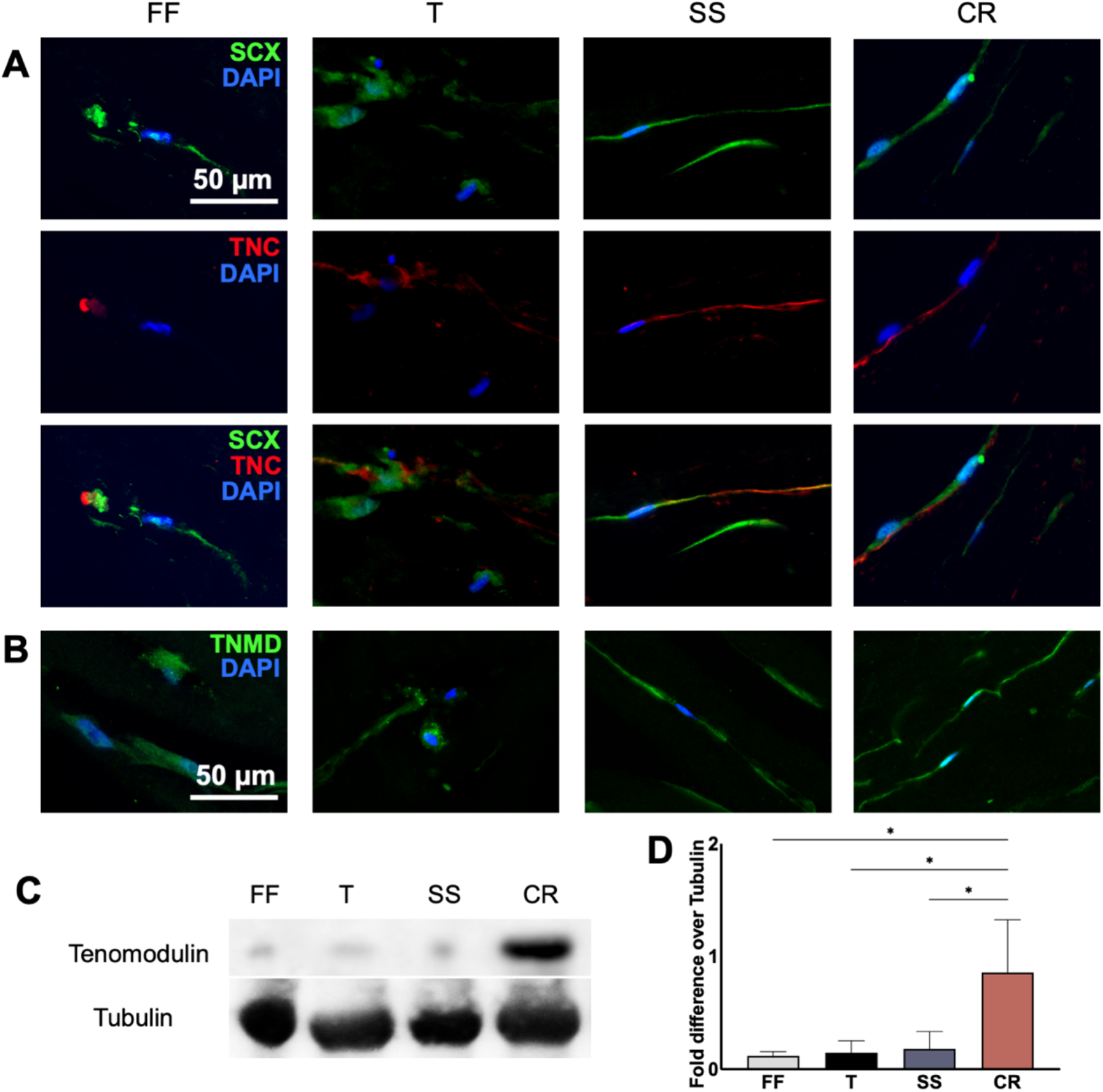
Tendon-specific markers of hBMMSC-seeded ADCs at day 21, after 48 h mechanoactivation and 19-day maturation of 20% static strain (SS) and 20% cyclic rest (CR) and their controls, free-floating (FF) and tethered (T). Representative images of immunofluorescent staining of (A) tenogenic transcription factor scleraxis (SCX) and tendon matrix protein tenascin-C (TNC), and (B) mature-tendon matrix protein tenomodulin (TNMD). (C) Western blot protein analysis of tenomodulin in the ADC hydrogels and (D) densitometry analysis of tenomodulin. Error bars ± SD and *p < 0.05, **p < 0.01 and ***p < 0.001 as per one-way ANOVA post-hoc multiple comparison analysis. Scale bars = 50 *μ*m.

## 4. Discussion

This study has shown that the GAE method rapidly fabricated ADC hydrogels that mimicked immature tendons both in composition and in stiffness [29,30]. The hydrogels were compliant under short-term (48 h) and high-strain uniaxial tensile loading, which not only increased the mechanical properties of the scaffolds but also committed seeded hBMMSCs to tenogenic differentiation in serum- and growth factor-free medium. The further 19-day maturation period of these mechanically activated hBMMSCs in ADC hydrogels in the absence of continued external mechanical stimulation showed cellular alignment and tendon-like matrix formation, highlighting the potential for short-term mechanoactivation driven tenogenic differentiation of MSCs.

It is widely accepted that scaffolds should exhibit characteristics of the intended tissue for optimized regeneration, preferably showing biomimicry in composition, structure, and mechanical properties. Tendons are predominantly comprised of collagen type I fibrils that are aligned to form anisotropic tissues resistant to high mechanical strains. In under 2 hours, GAE fabricates cell-seeded ADC hydrogels with tunable fibrillar content and alignment from acid-solubilized collagen type I [26,48]. The as made ADC scaffolds studied here were between 5 and 13 wt.% collagen, thereby approaching those of the native tendon range of approximately 18 – 38 wt.% collagen [22,61]. These values are significantly improved from traditional collagen hydrogels, as commercially available collagen type I in acidic solutions range between 2 to 10 mg/mL resulting in their highly hydrated nature with < 1 wt.% collagen [42,62]. These gels result in poor mechanical properties that are easily resorbed *in vivo* [24,62], and are unsuitable as force-transmitting scaffolds. Moreover, the collagen fibrils of these highly hydrated hydrogels are randomly oriented, limiting their potential use in the regeneration of anisotropic tissues, such as tendons. It has been observed that scaffolds with comparable stiffness with embryonic tendons promote tenogenic differentiation of seeded stem cells [7,20]. Previous studies on the traditional highly hydrated collagens showed that a six-week cell-driven remodeling period was required to induce fibrillar alignment and achieve a stiffness of approximately 1 MPa [25], *i.e.* in range of immature calf cruciate ligament modulus at 1-3 MPa [29] and neonatal chick tendon at 1-2 MPa [30]. However, by using the GAE method, these mechanical properties could be observed immediately post-fabrication or after the short-term (48 h) loading period depending on the needle diameter used. For example, 12G hydrogels (AM of 0.58 ± 0.14 MPa) reached 1.72 ± 0.49 and 1.43 ± 0.79 MPa after 20% static and cyclic loading, respectively. Interestingly, the mechanical stimulation profiles did not significantly affect the alignment of ADCs but appeared to increase their fibrillar thickness. This increase may be due to fibril fusion as previously reported [40], but it may also suggest the formation of fibril bundles, which would bring the ADCs closer to the hierarchical structure of native tendons. Nevertheless, the correlation between increased tensile properties and fibril thickness were not linear suggesting potential other factors influencing the gel mechanical properties.

Acellular investigations revealed mechanical compliance of ADC hydrogels with 20% loading profiles. On the other hand, it has been shown that strain levels above 9% can cause tenocytes to lose their elongated morphology [63–65], undergo apoptosis, as well as induce catabolic effects and inflammatory markers [66–68]. In contrast, although MSCs have been stimulated up to 15% strain in 2D, the studies did not report cytotoxic responses [14,69]. Moreover, in 2D culture, both 8 and 12% cyclic straining drove tenogenic differentiation of MSCs, where the former was deemed to provide the most optimal mechanical microenvironment for tenogenesis [70]. The importance of strain selection has been further highlighted as 4% cyclic loading induces osteogenic differentiation [70,71]. The rationale for applying 20% mechanoactivating strain, was predominantly from a materials perspective, where compliance and enhanced mechanical properties were observed in randomly oriented, dense collagen hydrogels [40]. As native tendons fail below 20% strain [72], the application of these stimulation profiles would result in damage to the tissue. Conversely, the 12G ADC hydrogels failed at approximately 25%, leaving the 20% tensile straining within the elastic region. It was hypothesized that since the scaffold was undamaged, there would be minimal negative response from the cells. The ADCs fabricated with 12G was further investigated as it provided more reproducible scaffolds than with the 14G needles in the GAE system [26].

Indeed, no harmful effects were detected on the seeded cells throughout the culture period. We observed that the 48 h short-term and high-strain loading mechanically activated the hBMMSCs and induced their commitment to tenogenic lineages, compared to free-floating and tethered controls, which instead suggested osteogenic and chondrogenic differentiation. These two control groups were investigated to account for and counteract the cellular contraction that was previously observed [48]. Mechanoactivation was conducted in the absence of serum and growth factors to fully understand the cellular response to mechanical loading only. The 48 h application of 20% static and cyclic profiles upregulated scleraxis expression by over 12-fold and downregulated RUNX2 and aggrecan, indicating tenogenesis over osteo- or chondrogenesis. Morphologically, initial loading significantly polarized and elongated the cellular cytoskeleton and nucleus, while tethered MSCs controls were observed to elongate at a reduced rate whereas free-floating controls did not change their initially rounded morphology. The elongation and reorientation of MSCs due to cyclic loading has previously been attributed to tenogenesis [10,38,70]. Interestingly, expressions of free-floating controls were upregulated for scleraxis (2-fold), RUNX2 (8-fold) and aggrecan (11-fold), suggesting that the collagen hydrogels in its dense and anisotropic form can promote various differentiation pathways of MSCs, which can be manipulated and directed to the target tissue lineage. All treatment groups stained positive for alpha smooth muscle actin, an indicator of cell-driven contraction as well as matrix remodeling [54,55]. Previous investigation of MSCs seeded in randomly oriented dense collagen showed that mechanical stimulation induced the formation of alpha smooth muscle actin [56]. This suggests that the alignment of fibrils may enhance the contractile forces that are exerted by the seeded MSCs. Additionally, alpha smooth muscle actin has also been linked to osteogenesis [73], which may have driven the RUNX2 expression of the free-floating controls. Where there are no other stimuli, the ADCs may provide optimal microenvironment for osteogenic differentiation of MSCs. Indeed, previous investigations of MSC-seeded ADC hydrogels have shown osteogenic differentiation, in vitro (Kamranpour et al., 2016), and ectopic bone formation, when subcutaneously implanted [74]. On the other hand, simply tethering ADCs upregulated scleraxis by 18-fold, whereas RUNX2 was significantly downregulated and aggrecan was maintained at baseline. This suggests that providing a countering force to the inherent contractile forces exhibited by the MSCs may play a role in achieving tenogenic commitment.

After the initial 48 h mechanoactivation period in serum- and growth factor-free medium, the scaffolds were tethered and matured for a further 19-day period under basal medium supplemented with ascorbic acid. The aim was to investigate whether the cells could maintain their tenogenic commitment without further stimulation, which potentially could be replicated as an *in vivo* implant. Previous studies have shown that 10% cyclical loading of 2D-cultured MSCs for 48 h increased tenogenic gene expressions, which were maintained for a further 48 h in the absence of further mechanical stimuli [10]. Therefore, we tethered the scaffolds for a further 19 days in a custom mould. During this maturation phase, all conditions showed extensive cellular remodeling, especially with collagen type I formation. In loaded scaffolds, protein analysis indicated the formation of tendon-like matrix, with the robust detection of collagens type III and VI, scleraxis, tenascin-C and tenomodulin. Collagen type III is found in tendon matrices [61] whereas collagen type VI and scleraxis have been associated with tendon matrix development and maturation [31,57,75]. In addition, tenascin-C and tenomodulin are mechanoresponsive proteins that aid in tendon cell proliferation and matrix assembly [59,76,77]. Western blot analysis indicated that there was significantly more tenomodulin detected in the scaffolds loaded with 20% cyclic rest, suggesting dynamic loading is crucial in tenogenesis. It also suggests that an initial upregulation of tenogenic genes may not translate into mature tendon protein formation. Cellular alignment has been suggested as a tenogenic marker [78,79], where MSCs that had undergone 20% static strain showed the most alignment. These discrepancies show different merits in both static and dynamic loading that may benefit from further investigation.

The present study indicates that an initial 20% uniaxial loading phase of ADCs in the absence of serum and growth factors can mechanoactivate seeded MSCs to commit to a tenogenic lineage that is carried throughout a longer-term maturation phase. However, ADCs may inherently be an appropriate tendon scaffold. Anisotropy through aligned nanofibrillar structure of scaffolds has been highlighted as a key factor in achieving tenogenic differentiation [80,81] and high cell infiltration and cell matrix remodeling *in vivo* of acellular ADCs have been reported [74]. Recently, an automated GAE system was developed allowing rapid throughput and fabrication of ADCs with higher CFD and enhanced fibrillar alignment [48]. These advantages over the handheld GAE system used in this study may enhance the tenogenic commitment by providing ADCs with tailored mechanical properties and alignment. Furthermore, to promote tenogenic differentiation in ADCs, only mechanical loading was applied, without the supplementation of either serum or growth factors. There is a lack of consensus on the optimal growth factor combination required for the activation of tenogenesis, as their differentiation pathways are yet unclear. Additionally, the lack of supplemental growth factors may provide advantages for clinical translation. The tenogenic matrix formation during the maturation phase suggests that this may be replicable following implantation *in vivo*, where after a short stimulatory period the scaffolds could be implanted to form a tendon-like tissue and aid healing *in situ*. There is potential to mature the scaffolds *in vivo*, where the scaffold can be sutured in place within a subcutaneous nursery site or in an injury site. MSC-seeded randomly oriented dense collagen gels were implanted and investigated in a subcutaneous site, where the scaffold was sutured and cyclic tensile loading was applied *in situ*, which resulted in significant scaffold remodeling and matrix deposition [82]. To enable direct implantation of these gels within a tendon defect model, a modified Kessler’s suturing technique was introduced and was observed to be stable up to 12 weeks in a lapine model [83,84]. By 12 weeks in the defect model, both acellular and tendon fibroblast-seeded scaffolds showed integration of the scaffolds with the native tendon and had significantly increased in mechanical properties with indications of ongoing fibrillar alignment [83]. Furthermore, ongoing clinical trials have shown that lyophilized collagen scaffolds with platelet-rich plasma can be sutured to “bridge” torn anterior cruciate ligament that perform similarly to traditional autografts after two years [85]. This suggests that there is scope to apply this principle to the ADCs, where the alignment and hydrogel nature of the scaffold may further aid tendon or anterior cruciate ligament healing.

The data presented in this study has shown that mechanical stimuli alone can drive tenogenic differentiation. Mechanical loading has been suggested to activate mechanosensitive ion channels [71], however the relationship between strain and differentiation pathways are still unclear and warrants further investigation. The compliancy with high-strains may be due to the larger elastic region exhibited by the ADC hydrogels. The stiffness of the hydrogels can dictate the differentiation pathways as well as cell behaviour by influencing the cellular response through the cytoskeleton [15,86]. Moreover, mechanical stimuli has been observed to activate several mechanosensors including integrins, focal adhesion kinase and cadherins that may affect tenogenic differentiation of MSCs [87]. The recently introduced “active gel theory” suggests that cells are active and have a gel-like viscoelastic outer layer comprised of actin-myosin filaments that is maintained by a constant consumption of energy [88,89]. Therefore, the encompassing scaffold hydrogel properties may affect the mechanical stimuli that is sensed by the internal gel, or the seeded cells. To investigate and understand the relevance of this theory to tissue engineering and MSC differentiation, further experimental studies observing the cellular response to various scaffold stiffness and strain levels may be of interest.

## 5. Conclusions

This work has shown that the tenogenic commitment of MSCs can be induced by short-term and high-strain mechanical stimulation of ADC hydrogels. The limitations of traditional highly hydrated collagen hydrogels were overcome by the rapid densification and fibrillar realignment during the GAE fabrication process. The short-term application of both static and dynamic loading promoted tenogenic gene expression that was carried during a longer-term maturation phase, even without external mechanical stimuli, which resulted in extensive matrix remodeling and mature tendon-like matrix formation. These results demonstrate the potential for predetermination of MSC fate to promote matrix remodeling and tissue healing *in vitro* that may be translate into guided *in vivo* healing to provide an alternative graft substitute. Whilst the mechanisms are yet unclear, the application of high-strain uniaxial loading promoted tenogenic differentiation in ADC scaffolds, thereby indicating strong potential of ADC hydrogels and application of mechanical stimulation for tendon tissue engineering.

## Supporting information

SI_mouldstl

## 6. Acknowledgments

Funding from the Research Institute McGill University Health Center start-up funds, Réseau de recherche en santé buccodentaire et osseuse (RSBO), Canada NSERC, FRQNT and CFI are gratefully acknowledged. DHR is a FRQS Junior 2 Research Scholar and HP’s funding was partially supported by McGill Engineering Doctoral Award and the Lorne Trottier Engineering Graduate Fellowship.

## References

[1] James R, Kesturu G, Balian G, Chhabra AB. Tendon: biology, biomechanics, repair, growth factors, and evolving treatment options. J Hand Surg Am 2008;33:102–12.

[2] Wang T, Gardiner BS, Lin Z, Rubenson J, Kirk TB, Wang A, et al. Bioreactor design for tendon/ligament engineering. Tissue Eng Part B Rev 2013;19:133–46.

[3] Lim WL, Liau LL, Ng MH, Chowdhury SR, Law JX. Current Progress in Tendon and Ligament Tissue Engineering. Tissue Eng Regen Med 2019;16:549–71.

[4] Deniz G, Kose O, Tugay A, Guler F, Turan A. Fatty degeneration and atrophy of the rotator cuff muscles after arthroscopic repair: Does it improve, halt or deteriorate? Arch Orthop Trauma Surg 2014;134:985–90.

[5] Ciardulli MC, Marino L, Lamparelli EP, Guida M, Forsyth NR, Selleri C, et al. Doseresponse tendon-specific markers induction by growth differentiation factor-5 in human bone marrow and umbilical cord mesenchymal stem cells. Int J Mol Sci 2020;21:1–19.

[6] Govoni M, Berardi AC, Muscari C, Campardelli R, Bonafè F, Guarnieri C, et al. An Engineered Multiphase Three-Dimensional Microenvironment to Ensure the Controlled Delivery of Cyclic Strain and Human Growth Differentiation Factor 5 for the Tenogenic Commitment of Human Bone Marrow Mesenchymal Stem Cells. Tissue Eng - Part A 2017;23:811–22.

[7] Brown JP, Finley VG, Kuo CK. Embryonic mechanical and soluble cues regulate tendon progenitor cell gene expression as a function of developmental stage and anatomical origin. J Biomech 2014;47:214–22.

[8] Dai L, Hu X, Zhang X, Zhu J, Zhang J, Fu X, et al. Different tenogenic differentiation capacities of different mesenchymal stem cells in the presence of BMP-12. J Transl Med 2015;13:1–14.

[9] Yin Z, Guo J, Wu T, Chen X, Xu L, Lin S, et al. Stepwise Differentiation of Mesenchymal Stem Cells Augments Tendon-Like Tissue Formation and Defect Repair In Vivo. Stem Cells Transl Med 2016;5:1106–16.

[10] Chen YJ, Huang CH, Lee IC, Lee YT, Chen MH, Young TH. Effects of cyclic mechanical stretching on the mRNA expression of tendon/ ligament-related and osteoblast-specific genes in human mesenchymal stem cells. Connect Tissue Res 2008;49:7–14.

[11] Kuo CK, Tuan RS. Mechanoactive tenogenic differentiation of human mesenchymal stem cells. Tissue Eng - Part A 2008;14:1615–27.

[12] Scott A, Danielson P, Abraham T, Fong G, Sampaio A V., Underhill TM. Mechanical force modulates scleraxis expression in bioartificial tendons. J Musculoskelet Neuronal Interact 2011;11:124–32.

[13] Juncosa-Melvin N, Boivin GP, Galloway MT, Gooch C, West JR, Sklenka AM, et al. Effects of cell-to-collagen ratio in mesenchymal stem cell-seeded implants on tendon repair biomechanics and histology. Tissue Eng 2005;11:448–57.

[14] Morita Y, Watanabe S, Ju Y, Xu B. Determination of optimal cyclic Uniaxial stretches for stem cell-to-tenocyte differentiation under a wide range of mechanical stretch conditions by evaluating gene expression and protein synthesis levels. Acta Bioeng Biomech 2013;15:71–9.

[15] Engler AJ, Sen S, Sweeney HL, Discher DE. Matrix Elasticity Directs Stem Cell Lineage Specification. Cell 2006;126:677–89.

[16] Rowlands AS, George PA, Cooper-White JJ. Directing osteogenic and myogenic differentiation of MSCs: Interplay of stiffness and adhesive ligand presentation. Am J Physiol - Cell Physiol 2008;295:1037–44.

[17] Islam A, Younesi M, Mbimba T, Akkus O. Collagen Substrate Stiffness Anisotropy Affects Cellular Elongation, Nuclear Shape, and Stem Cell Fate toward Anisotropic Tissue Lineage. Adv Healthc Mater 2016;5:2237–47.

[18] Murphy CM, Matsiko A, Haugh MG, Gleeson JP, O’Brien FJ. Mesenchymal stem cell fate is regulated by the composition and mechanical properties of collagen-glycosaminoglycan scaffolds. J Mech Behav Biomed Mater 2012;11:53–62.

[19] Park JS, Chu JS, Tsou AD, Diop R, Tang Z, Wang A, et al. The effect of matrix stiffness on the differentiation of mesenchymal stem cells in response to TGF-β. Biomaterials 2011;32:3921–30.

[20] Glass ZA, Schiele NR, Kuo CK. Informing tendon tissue engineering with embryonic development. J Biomech 2014;47:1964–8.

[21] Marturano JE, Schiele NR, Schiller ZA, Galassi T V., Stoppato M, Kuo CK. Embryonically inspired scaffolds regulate tenogenically differentiating cells. J Biomech 2016;49:3281–8.

[22] Kjær M. Role of Extracellular Matrix in Adaptation of Tendon and Skeletal Muscle to Mechanical Loading. Physiol Rev 2004;84:649–98.

[23] Thorpe CT, Birch HL, Clegg PD, Screen HRC. The role of the non-collagenous matrix in tendon function. Int J Exp Pathol 2013;94:248–59.

[24] Habibovic P, Bassett DC, Doillon CJ, Gerard C, McKee MD, Barralet JE. Collagen biomineralization in vivo by sustained release of inorganic phosphate ions. Adv Mater 2010;22:1858–62.

[25] Puetzer JL, Ma T, Sallent I, Gelmi A, Stevens MM. Driving hierarchical collagen fiber formation for functional tendon, ligament, and meniscus replacement. Biomaterials 2020:120527.

[26] Kamranpour NO, Miri AK, James-Bhasin M, Nazhat SN. A gel aspiration-ejection system for the controlled production and delivery of injectable dense collagen scaffolds. Biofabrication 2016;8:15018.

[27] Marelli B, Ghezzi CE, James-Bhasin M, Nazhat SN. Fabrication of injectable, cellular, anisotropic collagen tissue equivalents with modular fibrillar densities. Biomaterials 2015;37:183–93.

[28] Muangsanit P, Day A, Dimiou S, Ataç AF, Kayal C, Park H, et al. Rapidly formed stable and aligned dense collagen gels seeded with Schwann cells support peripheral nerve regeneration. J Neural Eng 2020;17:046036.

[29] Eleswarapu S V., Responte DJ, Athanasiou KA. Tensile properties, collagen content, and crosslinks in connective tissues of the immature knee joint. PLoS One 2011;6:1–7.

[30] McBride DJ, Trelstad RL, Silver FH. Structural and mechanical assessment of developing chick tendon. Int J Biol Macromol 1988;10:194–200.

[31] Pryce BA, Watson SS, Murchison ND, Staverosky JA, Dünker N, Schweitzer R. Recruitment and maintenance of tendon progenitors by TGFΒ signaling are essential for tendon formation. Development 2009;136:1351–61.

[32] Brown JP, Galassi T V., Stoppato M, Schiele NR, Kuo CK. Comparative analysis of mesenchymal stem cell and embryonic tendon progenitor cell response to embryonic tendon biochemical and mechanical factors. Stem Cell Res Ther 2015;6:1–8.

[33] Kapacee Z, Yeung CYC, Lu Y, Crabtree D, Holmes DF, Kadler KE. Synthesis of embryonic tendon-like tissue by human marrow stromal/mesenchymal stem cells requires a three-dimensional environment and transforming growth factor β3. Matrix Biol 2010;29:668–77.

[34] Otabe K, Nakahara H, Hasegawa A, Matsukawa T, Ayabe F, Onizuka N, et al. Transcription factor mohawk controls tenogenic differentiation of bone marrow mesenchymal stem cells in vitro and in vivo. J Orthop Res 2015;33:1–8.

[35] Wang T, Chen P, Zheng M, Wang A, Lloyd D, Leys T, et al. In vitro loading models for tendon mechanobiology. J Orthop Res 2018;36:566–75.

[36] Huang Y, Zheng L, Gong X, Jia X, Song W, Liu M, et al. Effect of cyclic strain on cardiomyogenic differentiation of rat bone marrow derived mesenchymal stem cells. PLoS One 2012;7.

[37] Huang Y, Jia X, Bai K, Gong X, Fan Y. Effect of fluid shear stress on cardiomyogenic differentiation of rat bone marrow mesenchymal stem cells. Arch Med Res 2010;41:497–505.

[38] Jang J-Y, Lee SW, Park SH, Shin JW, Mun C, Kim S-H, et al. Combined Effects of Surface Morphology and Mechanical Straining Magnitudes on the Differentiation of Mesenchymal Stem Cells without Using Biochemical Reagents. J Biomed Biotechnol 2011;2011:1–9.

[39] Khayat G, Rosenzweig DH, Khavandgar Z, Li J, Murshed M, Quinn TM. Low-Frequency Mechanical Stimulation Modulates Osteogenic Differentiation of C2C12 Cells. ISRN Stem Cells 2013;2013:1–9.

[40] Cheema U, Chuo CB, Sarathchandra P, Nazhat SN, Brown RA. Engineering functional collagen scaffolds: Cyclical loading increases material strength and fibril aggregation. Adv Funct Mater 2007;17:2426–31.

[41] Choi KM, Seo YK, Yoon HH, Song KY, Kwon SY, Lee HS, et al. Effect of ascorbic acid on bone marrow-derived mesenchymal stem cell proliferation and differentiation. J Biosci Bioeng 2008;105:586–94.

[42] Ghezzi CE, Marelli B, Muja N, Nazhat SN. Immediate production of a tubular dense collagen construct with bioinspired mechanical properties. Acta Biomater 2012;8:1813–25.

[43] Hotaling NA, Bharti K, Kriel H, Simon CG. DiameterJ: A validated open source nanofiber diameter measurement tool. Biomaterials 2015;61:327–38.

[44] Hoemann CD. Molecular and biochemical assays of cartilage components. Methods Mol. Med., vol. 101, New Jersey: Humana Press; 2004, p. 127–56.

[45] Fairag R, Rosenzweig DH, Ramirez-Garcialuna JL, Weber MH, Haglund L. Three-dimensional printed polylactic acid scaffolds promote bone-like matrix deposition in vitro. ACS Appl Mater Interfaces 2019;11:15306–15.

[46] Rosenzweig DH, Carelli E, Steffen T, Jarzem P, Haglund L. 3D-printed ABS and PLA scaffolds for cartilage and nucleus pulposustissue regeneration. Int J Mol Sci 2015;16:15118–35.

[47] Livak KJ, Schmittgen TD. Analysis of relative gene expression data using real-time quantitative PCR and the 2-ΔΔCT method. Methods 2001;25:402–8.

[48] Griffanti G, Rezabeigi E, Li J, Murshed M, Nazhat SN. Rapid biofabrication of printable dense collagen bioinks of tunable properties. Adv Funct Mater 2020;30:1903874.

[49] Jo CH, Lim HJ, Yoon KS. Characterization of Tendon-Specific Markers in Various Human Tissues, Tenocytes and Mesenchymal Stem Cells. Tissue Eng Regen Med 2019;16:151–9.

[50] Gumucio JP, Schonk MM, Kharaz YA, Comerford E, Mendias CL. Scleraxis is required for the growth of adult tendons in response to mechanical loading. JCI Insight 2020;5:2020.03.18.997833.

[51] Dyment NA, Hagiwara Y, Matthews BG, Li Y, Kalajzic I, Rowe DW. Lineage tracing of resident tendon progenitor cells during growth and natural healing. PLoS One 2014;9.

[52] Karsenty G. Role of Cbfa1 in osteoblast differentiation and function. Semin Cell Dev Biol 2000;11:343–6.

[53] Huang GT-J, Gronthos S, Shi S. Mesenchymal Stem Cells Derived from Dental Tissues vs. Those from Other Sources: Their Biology and Role in Regenerative Medicine. J Dent Res 2009;88:792–806.

[54] Kinner B, Zaleskas JM, Spector M. Regulation of smooth muscle actin expression and contraction in adult human mesenchymal stem cells. Exp Cell Res 2002;278:72–83.

[55] Arora PD, McCulloch CAG. Dependence of collagen remodelling on α-smooth muscle actin expression by fibroblasts. J Cell Physiol 1994;159:161–75.

[56] Ghezzi CE, Marelli B, Donelli I, Alessandrino A, Freddi G, Nazhat SN. The role of physiological mechanical cues on mesenchymal stem cell differentiation in an airway tract-like dense collagen-silk fibroin construct. Biomaterials 2014;35:6236–47.

[57] Sardone F, Santi S, Tagliavini F, Traina F, Merlini L, Squarzoni S, et al. Collagen VI–NG2 axis in human tendon fibroblasts under conditions mimicking injury response. Matrix Biol 2016;55:90–105.

[58] Taye N, Karoulias SZ, Hubmacher D. The “other” 15–40%: The Role of Non-Collagenous Extracellular Matrix Proteins and Minor Collagens in Tendon. J Orthop Res 2020;38:23–35.

[59] Dex S, Alberton P, Willkomm L, Söllradl T, Bago S, Milz S, et al. Tenomodulin is Required for Tendon Endurance Running and Collagen I Fibril Adaptation to Mechanical Load. EBioMedicine 2017;20:240–54.

[60] Buschmann J, Meier Bürgisser G. Structure and function of tendon and ligament tissues. Biomech. Tendons Ligaments, Elsevier; 2017, p. 3–29.

[61] Aparecida de Aro A, de Campos Vidal B, Pimentel ER. Biochemical and anisotropical properties of tendons. Micron 2012;43:205–14.

[62] Griffanti G, Nazhat SN. Dense fibrillar collagen-based hydrogels as functional osteoid-mimicking scaffolds. Int Mater Rev 2020:1–20.

[63] Yao L, Bestwick CS, Bestwick LA, Maffulli N, Aspden RM. Phenotypic drift in human tenocyte culture. Tissue Eng 2006;12:1843–9.

[64] Jiang Y, Liu H, Li H, Wang F, Cheng K, Zhou G, et al. A proteomic analysis of engineered tendon formation under dynamic mechanical loading in vitro. Biomaterials 2011;32:4085–95.

[65] Hannafin JA, Arnoczky SP, Hoonjan A, Torzilli PA. Effect of stress deprivation and cyclic tensile loading on the material and morphologic properties of canine flexor digitorum profundus tendon: An in vitro study. J Orthop Res 1995;13:907–14.

[66] Yang G, Im HJ, Wang JHC. Repetitive mechanical stretching modulates IL-1β induced COX-2, MMP-1 expression, and PGE2 production in human patellar tendon fibroblasts. Gene 2005;363:166–72.

[67] Li Z, Yang G, Khan M, Stone D, Woo SLY, Wang JHC. Inflammatory Response of Human Tendon Fibroblasts, to Cyclic Mechanical Stretching. Am J Sports Med 2004;32:435–40.

[68] Juncosa-Melvin N, Matlin KS, Holdcraft RW, Nirmalanandhan VS, Butler DL. Mechanical stimulation increases collagen type I and collagen type III gene expression of stem cell-collagen sponge constructs for patellar tendon repair. Tissue Eng 2007;13:1219–26.

[69] Sumanasinghe RD, Pfeiler TW, Monteiro-Riviere NA, Loboa EG. Expression of proinflammatory cytokines by human Mesenchymal stem cells in response to cyclic tensile strain. J Cell Physiol 2009;219:77–83.

[70] Nam HY, Pingguan-Murphy B, Abbas AA, Merican AM, Kamarul T. Uniaxial Cyclic Tensile Stretching at 8% Strain Exclusively Promotes Tenogenic Differentiation of Human Bone Marrow-Derived Mesenchymal Stromal Cells. Stem Cells Int 2019;2019.

[71] Fernandez-Yague MA, Trotier A, Demir S, Abbah SA, Larrañaga A, Thirumaran A, et al. A Self-Powered Piezo-Bioelectric Device Regulates Tendon Repair-Associated Signaling Pathways through Modulation of Mechanosensitive Ion Channels. Adv Mater 2021.

[72] Butler DL, Kay MD, Stouffer DC. Comparison of material properties in fascicle-bone units from human patellar tendon and knee ligaments. J Biomech 1986;19:425–32.

[73] Kalajzic Z, Li H, Wang LP, Jiang X, Lamothe K, Adams DJ, et al. Use of an alpha-smooth muscle actin GFP reporter to identify an osteoprogenitor population. Bone 2008;43:501–10.

[74] Miri AK, Muja N, Kamranpour NO, Lepry WC, Boccaccini AR, Clarke SA, et al. Ectopic bone formation in rapidly fabricated acellular injectable dense collagen-Bioglass hybrid scaffolds via gel aspiration-ejection. Biomaterials 2016;85:128–41.

[75] Murchison ND, Price BA, Conner DA, Keene DR, Olson EN, Tabin CJ, et al. Regulation of tendon differentiation by scleraxis distinguishes force-transmitting tendons from muscle-anchoring tendons. Development 2007;134:2697–708.

[76] Midwood KS, Chiquet M, Tucker RP, Orend G. Tenascin-C at a glance. J Cell Sci 2016;129:4321–7.

[77] Buschmann J, Meier Bürgisser G. Mechanobiology of tendons and ligaments. Biomech. Tendons Ligaments, Elsevier; 2017, p. 63–80.

[78] Skutek M, Van Griensven M, Zeichen J, Brauer N, Bosch U. Cyclic mechanical stretching modulates secretion pattern of growth factors in human tendon fibroblasts. Eur J Appl Physiol 2001;86:48–52.

[79] Galloway MT, Lalley AL, Shearn JT. The Role of Mechanical Loading in Tendon Development, Maintenance, Injury, and Repair. J Bone Jt Surgery-American Vol 2013;95:1620–8.

[80] Kishore V, Bullock W, Sun X, Van Dyke WS, Akkus O. Tenogenic differentiation of human MSCs induced by the topography of electrochemically aligned collagen threads. Biomaterials 2012;33:2137–44.

[81] Wang T, Thien C, Wang C, Ni M, Gao J, Wang A, et al. 3D uniaxial mechanical stimulation induces tenogenic differentiation of tendon-derived stem cells through a PI3K/AKT signaling pathway. FASEB J 2018;32:4804–14.

[82] Mudera V, Morgan M, Cheema U, Nazhat SN, Brown R. Ultra-rapid engineered collagen constructs tested in an in vivo nursery site. J Tissue Eng Regen Med 2007;1:192–8.

[83] Sawadkar P, Sibbons P, Ahmed T, Bozec L, Mudera V. Engineering of a Functional Tendon Using Collagen As a Natural Polymer. ACS Biomater Sci Eng 2019.

[84] Sawadkar P, Alexander S, Tolk M, Wong J, McGrouther D, Bozec L, et al. Development of a Surgically Optimized Graft Insertion Suture Technique to Accommodate a Tissue-Engineered Tendon In Vivo. Biores Open Access 2013;2:327–35.

[85] Murray MM, Fleming BC, Badger GJ, Freiberger C, Henderson R, Barnett S, et al. Bridge-Enhanced Anterior Cruciate Ligament Repair Is Not Inferior to Autograft Anterior Cruciate Ligament Reconstruction at 2 Years: Results of a Prospective Randomized Clinical Trial. Am J Sports Med 2020;48:1305–15.

[86] Trichet L, Le Digabel J, Hawkins RJ, Vedula SRK, Gupta M, Ribrault C, et al. Evidence of a large-scale mechanosensing mechanism for cellular adaptation to substrate stiffness. Proc Natl Acad Sci U S A 2012;109:6933–8.

[87] Jaiswal D, Yousman L, Neary M, Fernschild E, Zolnoski B, Katebifar S, et al. Tendon tissue engineering: Biomechanical considerations. Biomed Mater 2020;15.

[88] Prost J, Jülicher F, Joanny JF. Active gel physics. Nat Phys 2015;11:111–7.

[89] Joanny JF, Prost J. Active gels as a description of the actin-myosin cytoskeleton. HFSP J 2009;3:94–104.

